# Inhibition of Calcineurin Results in Increased Susceptibility to Voriconazole in the White-Nose Syndrome Fungus *Pseudogymnoascus destructans*

**DOI:** 10.64898/2026.09.03.749185

**Authors:** Aidan Marshall, Rebecca Busch, Tatiana Boluarte, José M. Vargas-Muñiz

## Abstract

*Pseudogymnoascus destructans* is a psychrophilic ascomycete fungus that causes White-Nose Syndrome in North American hibernating bats. *P. destructans* invades the tissue present in bats’ noses and wing membranes, causing the characteristic white nose and cupping erosion in the wings. *P. destructans* exhibits characteristics of a hemibiotrophic fungal pathogen, in which *P. destructans* can invade the host cells without inducing damage. During this stage, *P. destructans* hyphae assume a morphology similar to fungal plant pathogens during the biotrophic stage of infection. However, the mechanism by which *P. destructans* regulates its morphology and virulence is not fully understood. The phosphatase calcineurin is a key regulator of fungal stress response and virulence and could play a role in *P. destructans* virulence. Calcineurin can be inhibited using FK506 or Cyclosporin A (CSA). FK506 interacts with FKBP12, while CSA interacts with Cyclosporin A, and the protein-drug complex then inhibits calcineurin activity. Here, we show that *P. destructans* is intrinsically resistant to one calcineurin inhibitor (FK506), possibly because *P. destructans* FKBP12 has a lysine instead of the conserved arginine at position 89, which has been shown to reduce the affinity of the FKBP12-FK506 complex for calcineurin. Using cyclosporin A, we found that calcineurin is important for fungal growth. Additionally, calcineurin inhibition increases voriconazole inhibition. In contrast, calcineurin inhibition was antagonistic to the anti-cell wall drug micafungin. Taken together, calcineurin’s role in fungal growth and response to voriconazole is conserved in *P. destructans* but may play a unique role in *P. destructans* response to micafungin.

## Importance

White-Nose Syndrome, caused by the psychrophilic fungus *P. destructans*, has decimated hibernating bat populations in North America. Although a critical need for novel treatments exists, development has lagged because we still do not fully understand the mechanisms that regulate *P. destructans* infection. Here, we explore the role of calcineurin, a key phosphatase that regulates fungal pathogenesis, in *P. destructans* growth and response to antifungal drugs. We found that *P. destructans* calcineurin is involved in fungal growth. Additionally, calcineurin inhibition led to increased susceptibility to the antifungal voriconazole. However, calcineurin inhibition reduced susceptibility to the anti-cell wall drug micafungin, which is the opposite phenotype observed in other fungal pathogens. These findings suggest that some of calcineurin’s roles are conserved in *P. destructans*; however, this psychrophilic fungus offers new biology to uncover.

### Observation

*Pseudogymnoascus destructans* is a psychrophilic ascomycete fungus that grows as a saprobe in cold, dark, and damp environments like caves. *P. destructans* also causes a disease in North American hibernating bats known as White-Nose Syndrome (1). This name refers to the characteristic presentation of a white mold on the nose of the bat, though it can also be found on the ears, skin, and wing membranes of bats (2). Researchers first documented this fungus in the northeastern USA in 2006, and it has since been identified throughout the country and in neighboring Mexico and Canada (3). The disease causes bats painful discomfort during dermal tissue replacement, prematurely rousing them from torpor (4). Unlike other dermatophytes, which infect only the skin surface, *P. destructans* can invade healthy skin (4). Additionally, *P. destructans* exhibits morphological changes during infection (5). When growing on the surface of the skin, *P. destructans* grows as true hyphae, while when growing in the diagnostic cupping erosions, *P. destructans* adopts a bulbous morphology reminiscent of the biotrophic stage of hemibiotrophic plant pathogens (2, 5). Consistent with a possible biotrophic stage, *P. destructans* can invade bat keratinocytes without damaging the cells and actively inhibits apoptosis (6). Nonetheless, how *P. destructans* regulates these processes remains unknown.

A major regulator of fungal pathogenesis and morphology is the phosphatase calcineurin, which consists of a smaller regulatory subunit and a larger catalytic subunit (7, 8). Calcineurin inhibition weakens virulence in animal and plant pathogens such as *Aspergillus fumigatus, Candida albicans, Cryptococcus neoformans, Mucor circinelloides, Magnaporthe oryzae*, and *Ustilago maydis* (9–14). Furthermore, calcineurin regulates resistance to clinically used antifungals, including triazoles and echinocandins (7, 15, 16). The immunosuppressants FK506 (Tacrolimus) and Cyclosporin A both inhibit calcineurin activity and have been commonly used to study calcineurin’s role in fungal biology (7, 8). Both drugs bind to a target protein, FKBP12 for FK506 and cyclophilin A for CSA, and the drug-protein complex then binds and inhibits calcineurin. Despite calcineurin’s conserved role in fungal virulence, it remains unclear how calcineurin contributes to *P. destructans* biology.

To determine the role of *P. destructans* calcineurin, we first performed a minimum effective concentration (MEC) assay to determine which concentration of FK506 or CSA inhibits calcineurin. Using this approach, we found that *P. destructans* appears intrinsically resistant to FK506 (Fig 1A). Because resistance to FK506 often results from mutations in the *fkbp12* gene, we sequenced the gene to identify any such mutations. However, we found no mutations in the *fkbp12* gene compared with the reference genome (data not shown). Next, we aligned the amino acid sequences of *P. destructans, A. fumigatus*, and human FKBP12 orthologs using Clustal Omega (17). Using this approach, we found that *P. destructans* FKBP12 has a predicted N-terminus extension that is not present in human or *A. fumigatus* FKBP12s (Fig. 1B). Additionally, *Pdfkbp12* had a Lysine at position 89 instead of the conserved Arginine. Mutation of this Arginine in human FKBP12 (R43K) does not impact the affinity of FKBP12 for FK506, but it reduced the affinity of the FKBP12-FK506 complex to calcineurin (18). To test if the *P. destructans fkbp12* gene would confer resistance to FK506, we generated an *A. fumigatus* in which its native *fkbp12* gene was replaced with the *P. destructans* ortholog (Δ*fkbp12::pdfkbp12*). The Δ*fkbp12::pdfkbp12* strain shows no growth defect under basal conditions; however, this strain is resistant to FK506 (Fig 1C). This suggests that *P. destructans* FKBP12 might still bind FK506; however, the FKBP12-FK506 complex may have low affinity for calcineurin and therefore fail to inhibit the phosphatase. Nonetheless, the cause of this resistance needs to be defined, as the Δ*fkbp12::pdfkbp12* strain expresses full-length *P. destructans* FKBP12, and other regions, such as the N-terminal extension, could contribute to resistance.

**Figure 1.**
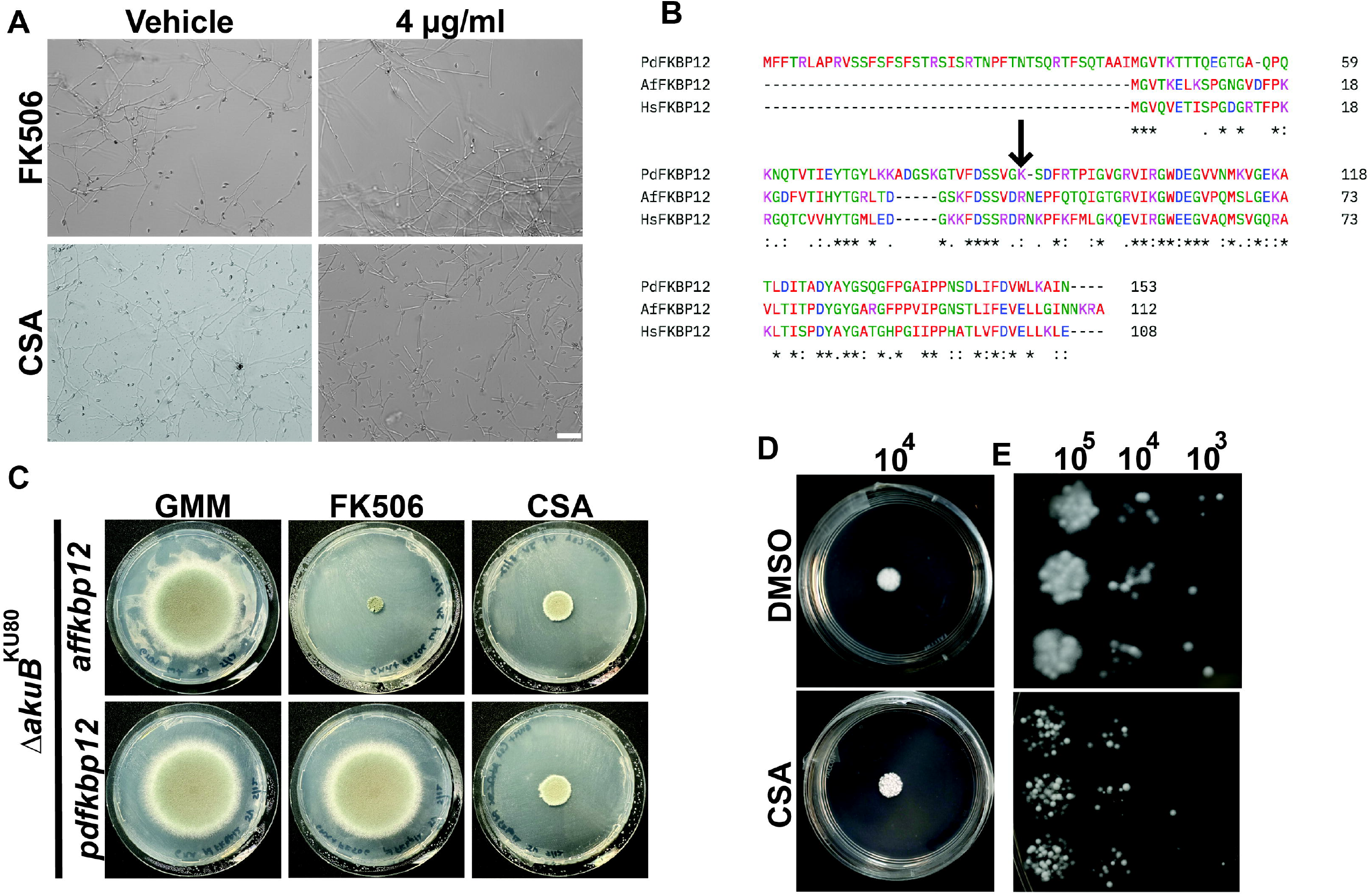
Inhibition of calcineurin by Cyclosporin A leads to reduced *P. destructans* growth. **(A)** Cyclosporin A (CSA) but not FK506 reduces hyphal growth. 2x10^4^ spores were inoculated into RPMI media containing DMSO (vehicle), FK506, or CSA and incubated for 72 hours at 14°C. Scale bar, 20µm. **(B)** *P. destructans, A. fumigatus*, and human FKBP12 protein sequence alignment show that *P. destructans* FKBP12 has an N-terminus extension and a Lysine at position 89 instead of a conserved arginine (arrow). Amino acid sequences were aligned using Clustal Omega. **(C)** Replacing the *A. fumigatus fkbp12* gene with the *P. destructans fkbp12* ortholog leads to FK506 resistance. 10^4^ spores were inoculated into GMM agar, GMM agar with FK506, or GMM agar with CSA, and incubated for 72 hours at 37°C. **(D-E)** GMM agar supplemented with 4 µg/ml CSA reduces *P. destructans* growth. **(D)** 10^4^ spores were inoculated in the center of the plate and incubated at 14°C for 15 days. **(E)** Minimum growth is observed when 10^3^ total spores are inoculated in GMM with CSA (4 µg/ml) and incubated for 15 days at 14°C.

*P. destructans* is susceptible to CSA, as we observed reduced growth starting at 4 µg/ml (Table S1). In the vehicle control, *P. destructans* grows robustly as mycelia in RPMI medium (Fig. 1A). However, 4 µg/ml CSA reduced growth in both RPMI liquid medium and a spot dilution assay on GMM agar (Fig. 1A, Fig 1D-E). As hyphal extension is a key phenotype for full virulence in filamentous fungi, it is possible that targeting calcineurin could reduce the virulence of *P. destructans*. However, the reduction in hyphal growth is less dramatic than in *A. fumigatus*, which may limit how effective CSA could be in controlling *P. destructans* infection (9). In addition to regulating hyphal morphology, calcineurin is important for fungal response to antifungals (7). For this reason, we tested whether CSA exposure increases susceptibility to the triazole voriconazole and the echinocandins micafungin and anidulafungin. *P. destructans* MEC was 0.25 µg/ml for voriconazole. Adding CSA reduced the MEC ∼16-fold to 0.015 µg/ml in RPMI (Table S1). *P. destructans* is capable of growing in voriconazole (0.5 µg/ml) in GMM agar but not in GMM agar supplemented with both CSA (4µg/ml) and voriconazole (0.5 µg/ml) (Fig 2A). This increased inhibition was also observed with the voriconazole e-strip, where GMM supplemented with CSA (4 µg/ml) showed a larger inhibition area than GMM alone (Fig. 2B). In contrast, calcineurin inhibition increases the MEC of micafungin from 4 µg/ml in RPMI to 8 µg/ml when RPMI is supplemented with CSA (4µg/ml) (Table S1). Similarly, the area of effect in the micafungin and anidulafungin e-strip test is reduced when CSA is supplemented to the media (Fig. 2B). This suggests a possible antagonistic interaction between CSA and the echinocandins. This is surprising, as calcineurin inhibition tends to be additive or synergistic with other antifungals, including echinocandins, in other fungal pathogens(7, 15). Taken together, calcineurin seems to regulate *P. destructans’* response to triazole, but not echinocandin. As genetic tools and new model systems become available, it will be possible to gain a more in-depth understanding of calcineurin’s role in *P. destructans* virulence and antifungal response.

**Figure 2.**
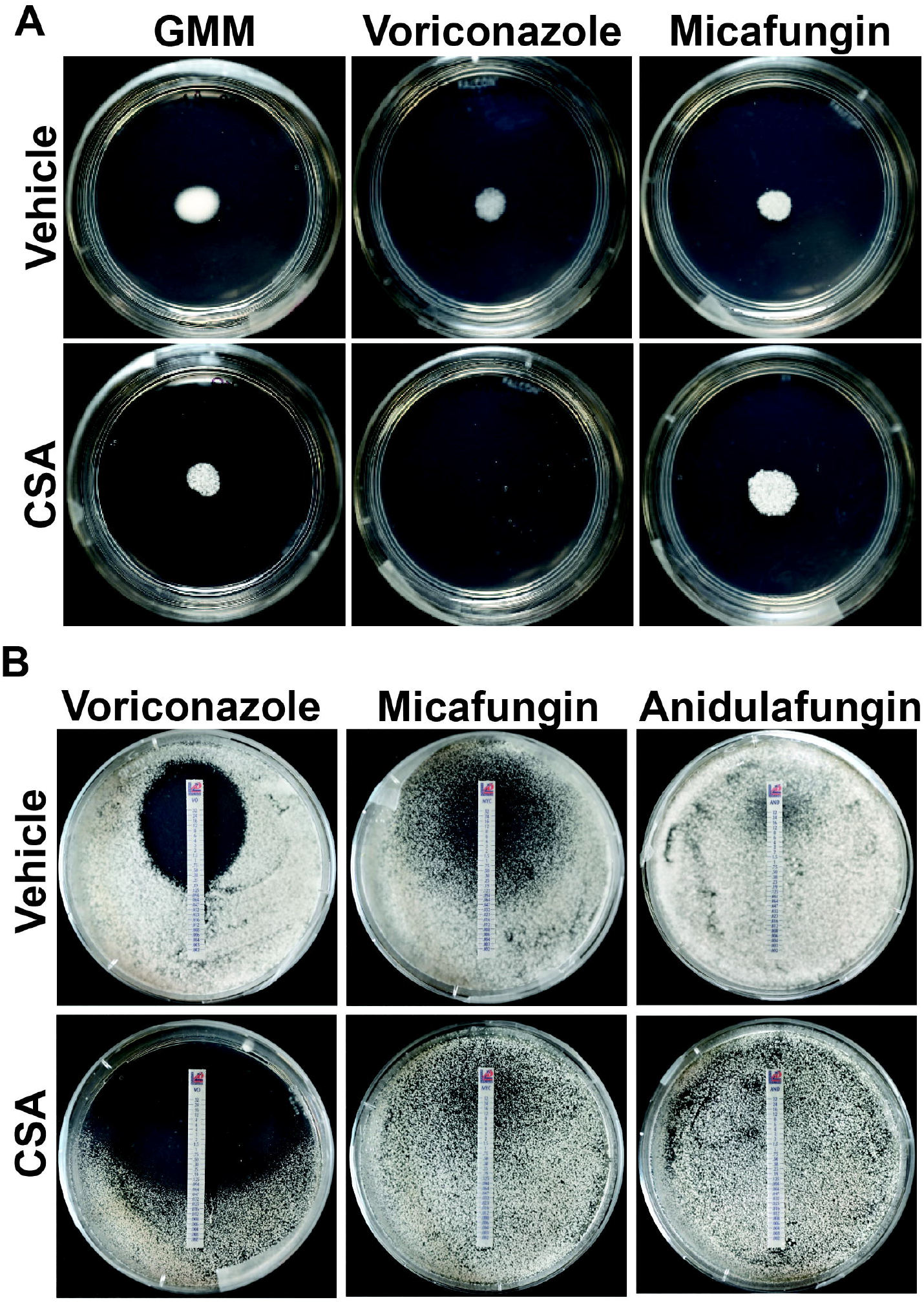
CSA exposure increases voriconazole inhibition. **(A)** *P. destructans* growth is fully inhibited when exposed to CSA (4µg/ml) and voriconazole (0.5 µg/µl). We inoculated 104 spores in the center of the plates and incubated them at 14°C for 15 days. **(B)** The voriconazole zone of inhibition increased when *P. destructans wa*s also exposed to CSA (4 µg/mL). 10^6^ spores were spread on GMM agar or GMM agar supplemented with CSA, and E-strips containing voriconazole, micafungin, or anidulafungin were placed in the center line of the plates. The plates were then incubated at 14°C for 15 days.

## Acknowledgments

This project is made possible through a grant from the National Fish and Wildlife Foundation (0406.23.81772), with support from the Bureau of Land Management (L20AC00505), U.S. Fish and Wildlife Service (F20AC00276), Avangrid Foundation, and Southern Company. R.J.B. was supported in part by the Virginia Tech Graduate Fellowship. This work was supported in part by the College of Science at Virginia Tech (Fund BR-COS1357) and made possible by funding from the Fralin Life Sciences Institute by FLSI Award 260027. The views and conclusions in this article are those of the authors and do not reflect the opinions or policies of the funding partners. We also want to thank members of the Vargas-Muñiz Lab for feedback on the manuscript.

## Figure Legends

**Table S1:**
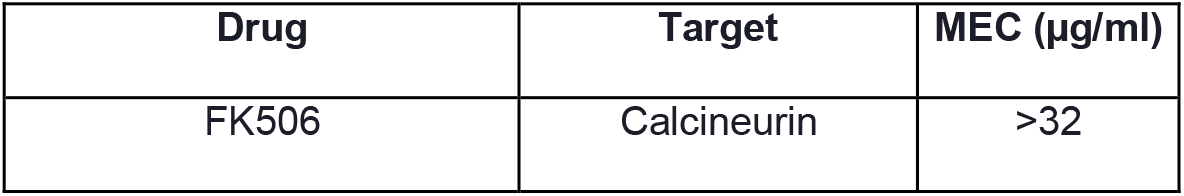
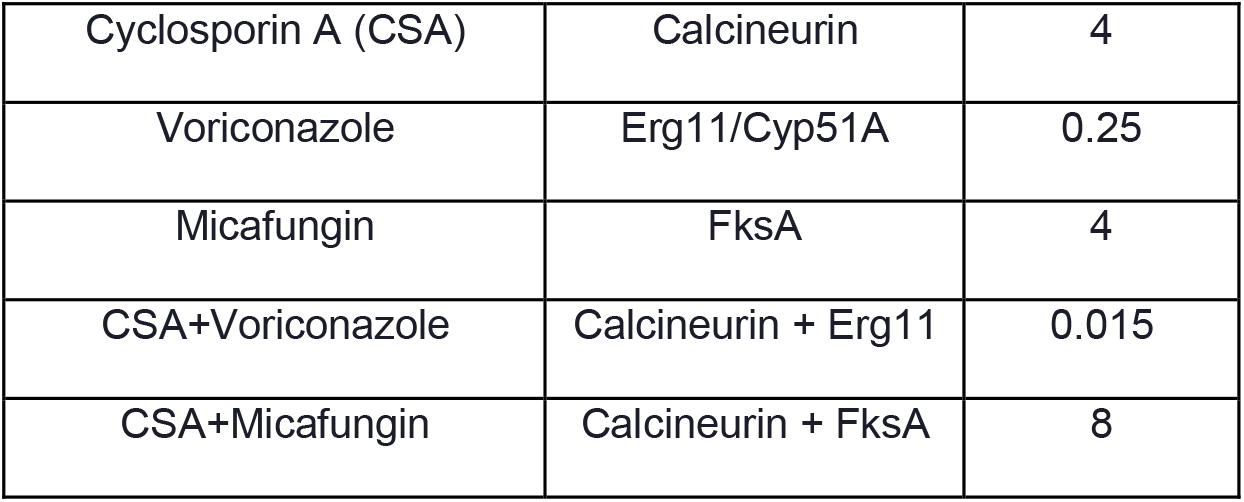
Minimum effective concentration (MEC) of drugs against *P. destructans*.

| Drug | Target | MEC (µg/ml) |
| --- | --- | --- |
| FK506 | Calcineurin | >32 |
| Cyclosporin A (CSA) | Calcineurin | 4 |
| Voriconazole | Erg11/Cyp51A | 0.25 |
| Micafungin | FksA | 4 |
| CSA+Voriconazole | Calcineurin + Erg11 | 0.015 |
| CSA+Micafungin | Calcineurin + FksA | 8 |

