## Supplementary material for "Inhibition of Calcineurin Results in Increased Susceptibility to Voriconazole in the White-Nose Syndrome Fungus *Pseudogymnoascus destructans*": Suplemental Materials and Methods

**Supplementary Materials and Methods**

***Strains, conditions, and media***

All experiments were conducted using the wild type *P. destructans* strain MYA-4855 (ATCC MYA-4855). The ∆*akuB*^KU80^ served as the wild type strain for *A. fumigatus*. *P. destructans* was grown on Glucose Minimal Media (GMM) at 14˚C unless otherwise specified, while *A. fumigatus* was grown in GMM at 37˚C.

***Generation of the ∆affkbp12::pdfkbp12* strain**

The *Aspergillus fumigatus* ∆*affkbp12::pdfkbp12* strain was generated by cloning the *A. fumigatus* terminator (898 bp) into the pUCGH plasmid in the SbfI and HindIII sites. The *Pdfkbp12* gene (GMDG_0425, 792bp) was amplified and fused to the *Affkbp12* promoter (~1 kb) using fusion PCR. The ~1.8 kb product was then digested with KpnI and NotI and cloned into the pUCGH vector containing the *Affkbp12* terminator. The resulting vector was verified by PCR and whole-plasmid sequencing. The vector was linearized, and the resulting 7 kb product was used to transform the ∆*akuB*^ku80^ strain. Successful transformants were selected for hygromycin resistance and then validated by PCR and Sanger sequencing of the genomic locus.

***Drug susceptibility assay***

For spot assays, 10^4^ *P. destructans* spores were inoculated onto GMM agar with Cyclosporin A (CSA, 4 µg/ml), CSA (4 µg/ml) and Voriconazole (250 or 500 ng/ml), CSA (4 µg/ml) and Caspofungin (1 µg/ml), or CSA (4 µg/ml and Micafungin (1 µg/ml). For the spore dilution assay, total spores ranging from 10^5^ to 10^2^ were plated on GMM agar with or without CSA (4 µg/ml) and an antifungal drug. Antifungal drugs used were Micafungin (1 µg/ml), and Voriconazole. Plates were incubated at 14˚C for 15 days. For E-strip assay, a suspension with 10^6^ spores in 500 µl was spread on GMM plates with CSA (4 µg/ml) or ethanol using sterile glass beads. After the spore solution dried, an E-strip containing Voriconazole, Micafungin, Caspofungin, or Anidulafungin was added to the plate. These plates were allowed to grow for 15 days at 14˚C before imaging. For the minimum effective concentration (MEC) assay, a total of 2.5 × 10^4^ spores were added to RPMI and RPMI+CSA (4 µg/ml) media containing different concentrations of micafungin or voriconazole in a 96-well plate. We incubated the plates for 72 hours at 14˚C before determining MEC values. Each assay was done in triplicate, and representative images were selected for each figure.
